# Gene modules associated with human diseases revealed by network analysis

**DOI:** 10.1101/598151

**Authors:** Shisong Ma, Jiazhen Gong, Wanzhu Zuo, Haiying Geng, Yu Zhang, Meng Wang, Ershang Han, Jing Peng, Yuzhou Wang, Yifan Wang, Yanyan Chen

**Author notes:** These authors contribute equally (Jiazhen Gong, Wanzhu Zuo).

## Abstract

Despite many genes associated with human diseases have been identified, disease mechanisms often remain elusive due to the lack of understanding how disease genes are connected functionally at pathways level. Within biological networks, disease genes likely map to modules whose identification facilitates etiology studies but remains challenging. We describe a systematic approach to identify disease-associated gene modules. A gene co-expression network based on the graphical Gaussian model (GGM) was constructed using the GTEx dataset and assembled into 652 gene modules. Screening these modules identified those with disease genes enrichment for obesity, cardiomyopathy, hypertension, and autism, which illuminated the molecular pathways underlying their pathogenesis. Using mammalian phenotypes derived from mouse models, potential disease candidate genes were identified from these modules. Also analyzed were epilepsy, schizophrenia, bipolar disorder, and depressive disorder, revealing shared and distinct disease modules among brain disorders. Thus, disease genes converge on modules within our GGM gene co-expression network, which provides a general framework to dissect genetic architecture of human diseases.

## INTRODUCTION

Human diseases often have genetic basis and their corresponding gene-disease associations are widely documented (1–4). However, many diseases, such as autism and other brain disorders, exhibit vast genetic heterogeneity, and their disease mechanisms remain elusive, despite a large number of disease genes have been identified (5,6). Growing evidences suggested that, within biological networks, disease genes usually do not distribute randomly but tend to map to modules that include subsets of genes functioning together in the same or similar pathways (7,8). Identifying such disease-associated gene modules will illuminate key molecular pathways underlying human diseases and help to bridge the knowledge gap between the genome and the clinic (6,9).

Systems biology methods have been developed to identify disease-associated gene modules. One approach is based on gene or protein co-expression analysis. Such analysis usually adopted the weighted gene co-expression network (WGCNA) method, which uses a power function of Pearson’s correlation coefficient to measure expression similarity (10). For example, Parikshak et al. constructed a WGCNA gene co-expression network using early brain developmental samples and identified gene modules enriched with autism genes (11). Huan et al. used whole blood transcriptomes from normal and hypertensive individuals to build WGCNA networks and found blood pressure-associated gene modules (12). Gandal et al. constructed co-expression network from cerebral cortex transcriptome across five major neuropsychiatric disorders using a robust WGCNA method and identified shared modules among these diseases (13). At the same time, Emilsson et al. also built a protein WGCNA co-regulatory network from serum proteome data, identified modules, and correlated the modules to cardiovascular and metabolic diseases (14). However, these network analyses were based on transcriptomes derived from single or limited tissue types, which usually produced large gene modules that could hinder subsequent functional analysis. Another approach is based on protein-protein interaction (PPI) network analysis. For example, O’Roak et al. identified *de novo* autism spectrum disorder risk genes and found 39% of them mapped to a highly interconnected PPI network (15). Li et al. clustered PPI network from BioGRID into modules and identified 2 of them enriched with autism genes (16). While Menche et al. also used PPI network to identify disease modules and reveal disease-disease relationship (8). On the other hand, a MAGI method was also proposed to integrate co-expression and PPI data to pinpoint autism-associated gene modules (17).

Although considerable progress have been made, disease gene modules have not been detected for many diseases. For those that have been analyzed, there is still much room for improvement. We describe a systematic approach to identify gene modules associated with human diseases. A human gene co-expression network was constructed based on the graphical Gaussian model (GGM) using the Genotype-Tissue Expression (GTEx) transcriptome dataset (18). GGM employs partial correlation coefficient to measure expression similarity between genes, which performs much better than the commonly used Pearson’s correlation coefficient in network construction (19,20). The network assembled into 652 gene co-expression modules. Screening these modules identified those with disease gene enrichment for obesity, cardiomyopathy, hypertension, autism, and other human diseases with major impacts on public health worldwide, elucidating the molecular pathways underlying their pathogenesis. Epilepsy, schizophrenia, bipolar disorder, and depressive disorder were also analyzed, revealing shared and distinct disease modules among brain disorders. Our approach provides a general framework to dissect genetic basis of human diseases.

## RESULTS

### GGM gene co-expression network and disease gene modules identification

We developed a systematic approach to identify gene modules associated with human diseases (Figure 1A). A human gene co-expression network based on the graphical Gaussian model was constructed using publicly available transcriptome data from the GTEx project (18). The project has created a resource of gene expression data from ‘normal’, non-diseased tissues, which supported a range of studies, including co-expression analysis (21,22). Our work used GTEx V7 release that contains transcriptome data for 11688 samples spanning 53 tissue types from 714 postmortem donors (18). The analysis followed a procedure we published previously to construct a GGM gene co-expression network across all 53 tissue types (23,24). The resulted network, HsGGM2019, contains 166980 co-expressed gene pairs among 18425 protein-coding genes (Table S1). Via the MCL algorithm (25), 652 gene co-expression modules with 9 or more genes were identified from the network. Considering some genes might participate in multiple pathways, the modules were expanded by including outside genes connected with >= 4 genes within the original modules. Accordingly, 14589 genes assembled into 652 modules containing 9 to 354 genes each, with 3733 genes belonging to multiple modules (Figure 1B and Table S2). The rest 3836 genes organized into smaller modules that were not considered hereafter.

**Figure 1.**
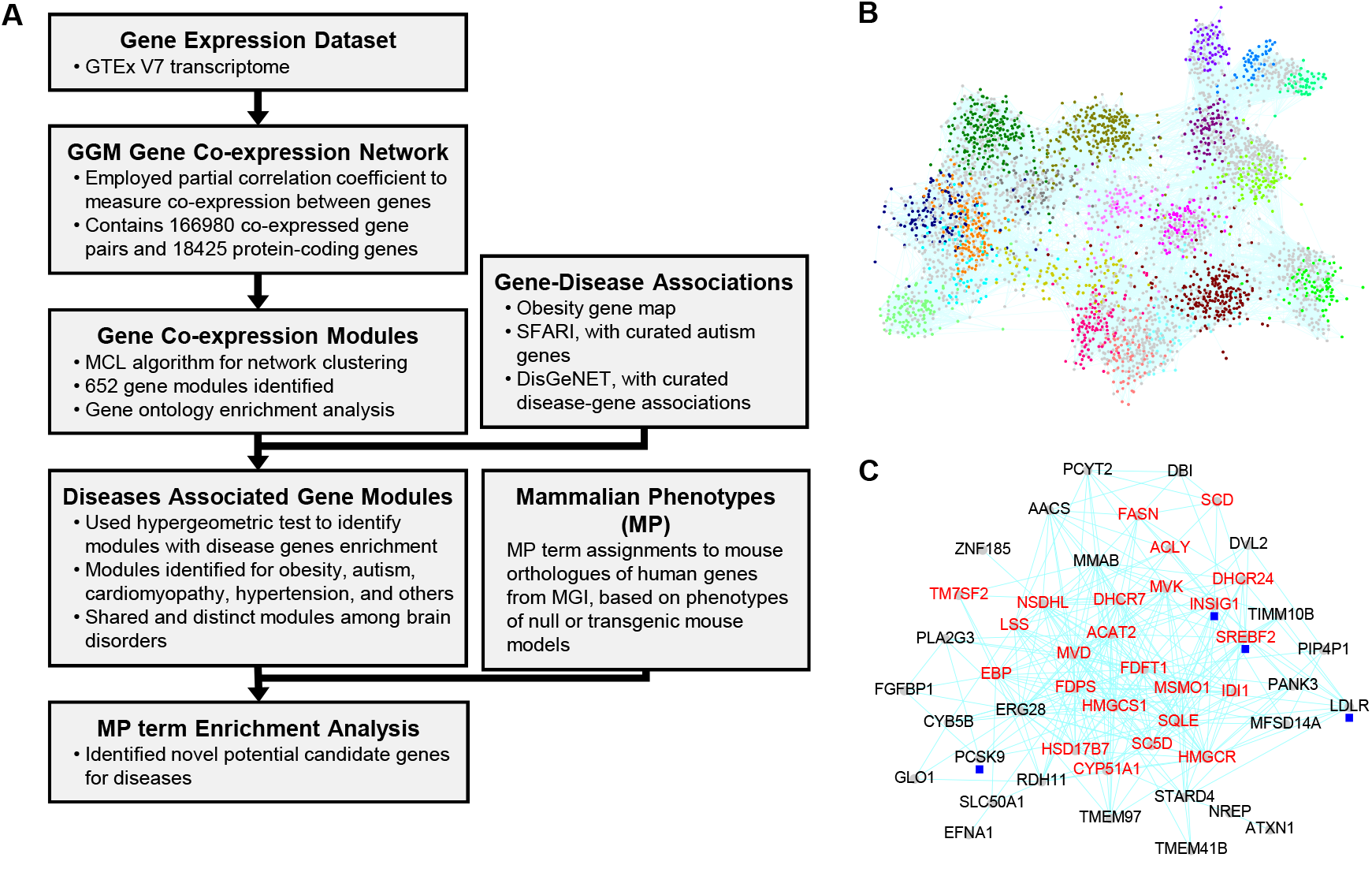
Experimental design and overview of the identified modules. **(A)** The analysis pipeline. **(B)** A sub-network for the largest 20 modules. Nodes represent genes and co-expressed genes are connected by edges. Colors of nodes indicate module identities, except that nodes in grey color are those belong to multiple modules. **(C)** A gene module (#71) for cholesterol biosynthesis. Highlighted in red are cholesterol biosynthesis genes. Labeled with blue square are genes encoding cholesterol homeostasis regulators.

The modules contain co-expressed genes that, according to guilt-by-association, might function in the same or similar pathways. Gene Ontology (GO) analysis identified 339 of them with enriched GO terms (Benjamini-Hochberg adjusted pValue [*P*]<=1E-2) (Table S3). Their biological relevance are exemplified by three modules involved in lipid biosynthesis, transport, and storage. Module #71, expressed broadly in various tissues (Figure S1), is enriched with 24 cholesterol biosynthesis genes (*P*=9.44E-45) (Figure 1C), including genes for enzymes catalyzing 23 of all 24 reaction steps for synthesizing cholesterol from acetyl-CoA (Figure S2). It also contains *LDLR*, *PCSK9*, *SREBF2*, and *INSIG1*, encoding components of the PCSK9-LDLR and SREBP-SCAP-Insig complexes that regulate cholesterol homeostasis (26). Module #30, containing genes with restricted expression towards liver (Figure S1), is enriched with high-density lipoprotein (HDL) particle genes (*P*=1.55E-13), such as *APOA1*, *APOA2*, and *APOA5*. This module could function in HDL-mediated reverse cholesterol transport. Module #18, expressed biasedly in fat (Figure S1), is enriched with lipid storage genes (*P*=6.96E-11). It contains *PPARG*, encoding a key transcription factor regulator of adipogenesis, and its target genes involved in adipocyte differentiation and metabolism, like *PLIN1*, *FABP4*, *LEP*, and *ADIPOQ* (27). Modules were also revealed for other processes, such as those functioning in specific organelles or tissues (#1, #27, #50), metabolism pathways (#54, #126, #288), immunity pathways (#6, #8, #15, #73), or general cellular pathways (#3, #28, #29) (see Table 1 and Table S3 for these modules’ enriched GO, same as below).

**Table 1.** Enriched GO terms for selected modules.

| M# | Enriched GO | P | M# | Enriched GO | P |
| --- | --- | --- | --- | --- | --- |
| 1 | cilium | 1.61E-61 | 83 | MHC class I protein complex | 7.55E-15 |
| 2 | mitochondrion | 3.8E-185 | 85 | muscle contraction | 1.47E-25 |
| 3 | cell cycle | 1.5E-146 | 92 | muscle contraction | 1.96E-36 |
| 6 | immune response | 1.22E-70 | 93 | synaptic signaling | 4.13E-06 |
| 7 | chromosome organization | 6.21E-31 | 95 | inflammatory response | 6.46E-19 |
| 8 | immune response | 8.46E-73 | 101 | extracellular matrix | 1.72E-36 |
| 9 | neurogenesis | 4.31E-18 | 105 | proteolysis | 2.15E-07 |
| 11 | synaptic signaling | 3.7E-36 | 113 | G protein-coupled receptor activity | 2.23E-04 |
| 14 | myelination | 6.52E-10 | 114 | visual perception | 2.29E-16 |
| 15 | adaptive immune response | 4.30E-75 | 124 | synaptic signaling | 7.96E-07 |
| 18 | lipid storage | 6.96E-11 | 126 | cGMP metabolic process | 9.22E-04 |
| 20 | synaptic signaling | 5.92E-21 | 131 | extracellular matrix | 3.59E-12 |
| 22 | mitochondrion | 2.40E-37 | 146 | synaptic signaling | 7.33E-10 |
| 27 | lysosome | 1.13E-43 | 154 | alveolar lamellar body | 4.34E-08 |
| 28 | ribosome biogenesis | 1.30E-63 | 168 | heat shock protein binding | 2.53E-02 |
| 29 | protein folding | 9.60E-35 | 174 | regulation of hormone levels | 8.82E-07 |
| 30 | high-density lipoprotein particle | 1.55E-13 | 186 | cell-cell adhesion | 1.57E-16 |
| 34 | regulation of cell death | 7.43E-17 | 187 | synaptic signaling | 5.26E-05 |
| 35 | synaptic signaling | 8.37E-17 | 191 | organic acid metabolic process | 1.77E-19 |
| 39 | muscle contraction | 6.72E-29 | 197 | behavior | 1.87E-06 |
| 50 | brush border | 1.19E-13 | 229 | regulation of synaptic plasticity | 9.02E-03 |
| 51 | thyroid hormone generation | 2.02E-08 | 231 | circadian regulation of gene expression | 8.27E-07 |
| 54 | glucocorticoid metabolic process | 1.10E-09 | 249 | neurotransmitter binding | 9.66E-03 |
| 56 | muscle structure development | 7.37E-29 | 264 | MHC class II protein complex | 6.70E-31 |
| 58 | muscle contraction | 5.56E-38 | 277 | neuropeptide signaling pathway | 9.62E-04 |
| 60 | apoptotic process | 2.58E-07 | 281 | ion transmembrane transporter activity | 1.02E-06 |
| 63 | vasculature development | 3.31E-18 | 288 | dopamine metabolic process | 2.08E-08 |
| 66 | extracellular matrix | 1.59E-75 | 305 | sodium ion homeostasis | 3.32E-03 |
| 67 | cilium | 1.14E-22 | 358 | neuron part | 4.31E-03 |
| 70 | fatty acid oxidation | 6.56E-31 | 374 | mitochondrial ATP synthesis coupled electron transport | 2.19E-20 |
| 71 | cholesterol biosynthetic process | 9.44E-45 | 396 | regulation of vascular smooth muscle cell differentiation | 9.31E-03 |
| 72 | anion transmembrane transporter activity | 3.25E-16 | 442 | inward rectifying potassium channel | 1.49E-03 |
| 73 | immune response | 2.77E-40 | 466 | synapse | 5.20E-03 |
| 76 | synaptic signaling | 7.64E-20 | 520 | fatty acid biosynthetic process | 1.12E-08 |
| 77 | nervous system development | 1.07E-05 | 546 | feeding behavior | 6.44E-04 |
| 82 | rhythmic process | 3.49E-10 | 598 | forebrain neuron differentiation | 1.76E-09 |

M#, module ids; P, BH-adjusted pValue for GO enrichment.

The modules were then screened for association with diseases. A module is considered as disease-associated if it has disease genes enrichment (permutation-based False Discovery Rate [FDR]<=0.05). Curated gene-disease associations for obesity, autism, and other diseases, retrieved from obesity gene map, SFARI (2019/01/15 update), and DisGeNET (v5.0) (3-5), were queried against the modules. Gene modules associated with obesity, cardiomyopathy, hypertension, autism and other human diseases were identified (Table 2 and Table S4), as discussed below in details.

**Table 2.** Gene modules associated with obesity, cardiomyopathy, hypertension, and autism.

| M# | Disease<br>gene<br>counts | OR | BH<br>Adjusted<br>pValue | FDR |
| --- | --- | --- | --- | --- |
| Obesity |  |  |  |  |
| 18 | 22 | 10.5 | 8.26E-14 | <0.0001 |
| 288 | 6 | 21.1 | 2.06E-05 | 5E-05 |
| 30 | 13 | 5.1 | 1.38E-04 | 0.0002 |
| 60 | 9 | 6.9 | 3.18E-04 | 0.0003 |
| 101 | 10 | 6.0 | 3.18E-04 | 0.0003 |
| 34 | 12 | 4.4 | 7.19E-04 | 0.0007 |
| 546 | 4 | 21.0 | 7.19E-04 | 0.001 |
| 50 | 9 | 5.2 | 1.71E-03 | 0.0024 |
| 277 | 4 | 13.1 | 4.42E-03 | 0.0048 |
| 174 | 5 | 7.3 | 1.16E-02 | 0.0137 |
| 396 | 3 | 13.1 | 2.42E-02 | 0.0369 |
| 191 | 5 | 5.8 | 2.71E-02 | 0.0373 |
| Dilated Cardiomyopathy (DCM) |  |  |  |  |
| 85 | 16 | 64.5 | 4.49E-22 | <0.0001 |
| 92 | 11 | 33.9 | 2.60E-12 | <0.0001 |
| 70 | 6 | 25.3 | 3.63E-06 | <0.0001 |
| 39 | 8 | 12.6 | 6.56E-06 | <0.0001 |
| 58 | 4 | 10.5 | 7.48E-03 | 0.0233 |
| Hypertrophic Cardiomyopathy (HCM) |  |  |  |  |
| 2 | 22 | 16.5 | 7.50E-17 | <0.0001 |
| 85 | 10 | 32.8 | 3.68E-11 | <0.0001 |
| 70 | 7 | 26.7 | 2.19E-07 | 3E-05 |
| 92 | 5 | 12.7 | 7.64E-04 | 0.0024 |
| 39 | 5 | 6.8 | 1.07E-02 | 0.0451 |
| Hypertensive Disease (Hypertension) |  |  |  |  |
| 374 | 7 | 35.9 | 4.96E-08 | <0.0001 |
| 95 | 8 | 10.7 | 4.92E-05 | <0.0001 |
| 39 | 13 | 5.5 | 4.92E-05 | <0.0001 |
| 30 | 12 | 6.0 | 4.92E-05 | 3E-05 |
| 281 | 5 | 16.6 | 3.33E-04 | 0.0003 |
| 66 | 9 | 6.5 | 3.33E-04 | 0.0004 |
| 305 | 4 | 17.6 | 1.55E-03 | 0.0016 |
| 34 | 10 | 4.7 | 1.64E-03 | 0.0016 |
| 54 | 6 | 7.4 | 3.21E-03 | 0.0026 |
| 63 | 6 | 5.8 | 9.47E-03 | 0.0147 |
| 126 | 4 | 10.1 | 9.47E-03 | 0.0147 |
| 101 | 6 | 4.5 | 3.43E-02 | 0.0421 |
| 105 | 4 | 6.9 | 3.43E-02 | 0.0421 |
| Autism |  |  |  |  |
| 7 | 48 | 5.9 | 2.41E-20 | <0.0001 |
| 186 | 17 | 18.7 | 3.42E-17 | <0.0001 |
| 77 | 14 | 5.8 | 9.88E-06 | <0.0001 |
| 35 | 16 | 3.8 | 3.89E-04 | 0.0003 |
| 20 | 18 | 3.3 | 6.56E-04 | 0.0006 |
| 11 | 23 | 2.8 | 6.97E-04 | 0.0006 |
| 168 | 7 | 7.3 | 1.43E-03 | 0.0015 |
| 9 | 15 | 2.5 | 3.91E-02 | 0.0356 |
| 358 | 4 | 8.0 | 4.04E-02 | 0.04 |
| 325 | 4 | 7.4 | 4.93E-02 | 0.0461 |
| 131 | 6 | 4.4 | 5.17E-02 | 0.0429 |
| 146 | 6 | 4.4 | 5.17E-02 | 0.0429 |

M#, module id. OR, odd ratio. FDR, false discovery rate.

### Gene modules associated with obesity

As an example, Module #18 for lipid storage was found to be associated with obesity (Table 2), a disease involving excessive body fat that affects 12% adults worldwide (28). Besides environment, genetic factors influence obesity susceptibility. According to obesity gene map, 22 out of the 115 genes within Module #18 are obesity genes (Figure 2A), including *ADIPOQ* and *LEP*, whose genetic polymorphisms are risk factors for obesity (29). Comparing to the whole network, obesity genes are 10.5 fold enriched (Odd Ratio, OR) within the module (FDR<0.0001). We examined phenotypes of transgenic or null mouse models developed for the genes within the module, using Mammalian Phenotype (MP) Ontology assignments from MGI (30), to investigate if other genes also relate to obesity. The MP *abnormal body weight* is associated with 33 genes and significantly enriched within the module (*P*=4.91E-07) (Figure 2A, see Table S5 for MP enrichment analysis results for all modules, and refer to MATERIALS AND METHODS for analysis details). Among them are 15 obesity genes, and the rest 18 might represent additional candidate genes. Note that these candidate genes derived from mouse models need to be further verified by, i.e., human population genetics studies. Additionally, Module #18 contains 7 genes for insulin resistance (OR=17.4, FDR=0.0001), a syndrome that could result from obesity (31). It also has 13 genes with the MP *insulin resistance* (*P*=2.23E-10), again expanding the disease’s candidate gene list.

**Figure 2.**
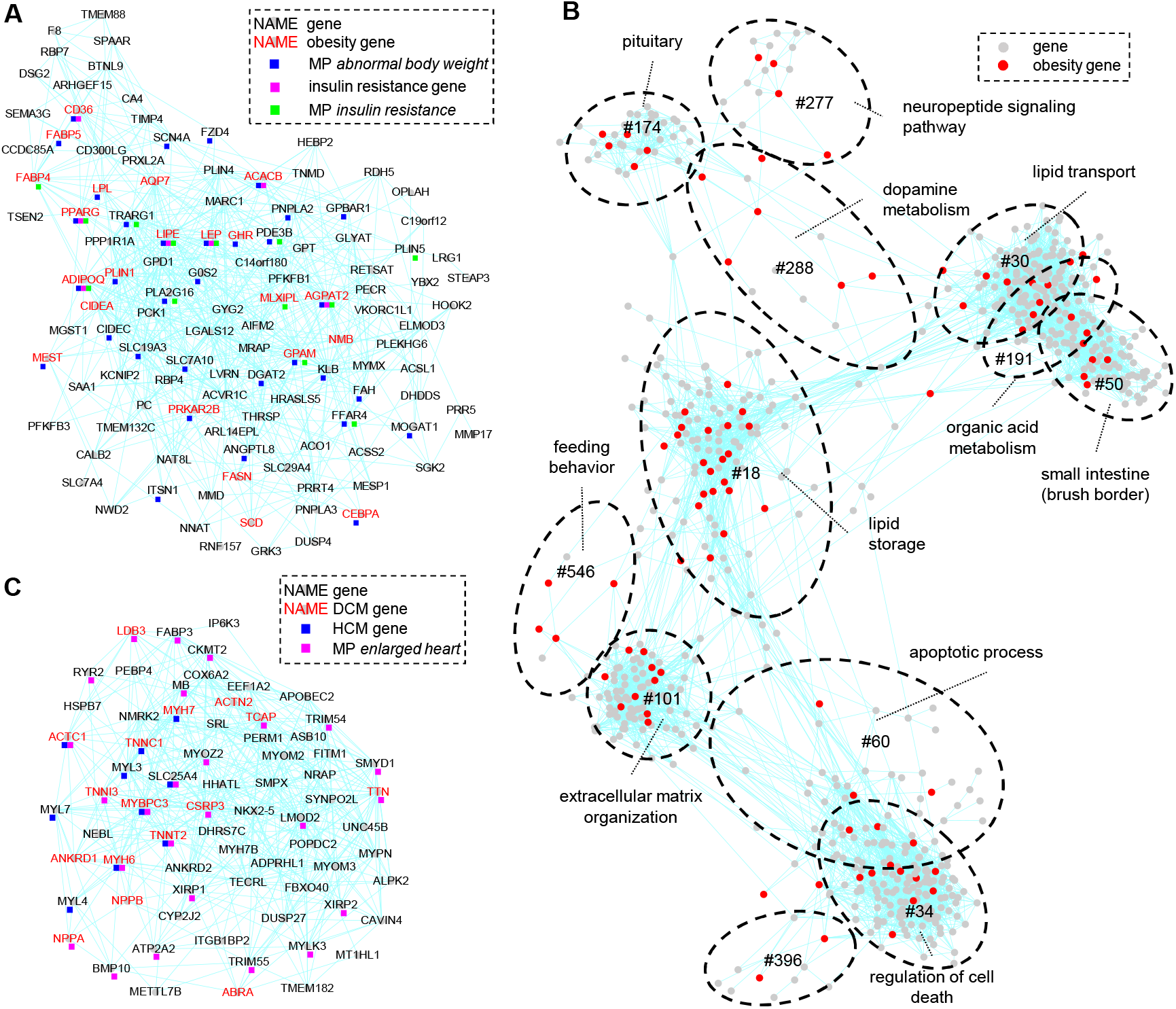
Gene modules associated with obesity and cardiomyopathy. **(A)** A module (#18) associated with obesity. Obesity genes, insulin resistance genes, and genes with MPs *abnormal body weight* and *insulin resistance* are indicated. **(B)** A sub-network including all modules associated with obesity. Dashed circles outline the approximate positions of the modules. **(C)** A module (#85) associated with cardiomyopathy. Disease genes for dilated cardiomyopathy (DCM) and hypertrophic cardiomyopathy (HCM) are indicated.

Also discovered are 11 additional obesity modules (Figure 2B), 10 of which function in dopamine metabolism (#288), neuropeptide signaling pathway (#277), feeding behavior regulation (#546), lipid transport (#30), organic acid metabolism (#191), extracellular matrix organization (#101), regulation of cell death (#34), and apoptotic process (#60), or have biased expression in small intestine (#50) and pituitary (#174), respectively (Figure S3). Among them, dopamine relates to obesity via modulating appetite, while within Module #277 and #174 are also genes regulating feeding behavior, such as *HCRT*, *PMCH*, and *POMC* (32–34).

### Gene modules associated with cardiomyopathy

Modules were identified for diseases affecting specific tissues or organs. Cardiomyopathy is a heart muscle disease, where the heart muscle becomes enlarged, thick or stiff (35). Dilated cardiomyopathy (DCM) and hypertrophic cardiomyopathy (HCM) are among the major types of cardiomyopathy. Six gene modules are associated with DCM and/or HCM (Table 2). Three DCM modules, #85, #92, #39 (OR>=12.6, FDR<0.0001), have enriched GO *muscle contraction* (*P*<=1.47E-25), indicating their muscle-related functions. While Module #85 and #92 are specifically expressed in heart and skeletal muscles, #39 is also expressed in tissues with smooth muscles like artery, colon, and esophagus (Figure S4). Pathway analysis (36) indicated that Module #85 and #92 function in striated muscle contraction and #39 in smooth muscle contraction. DCM is also associated with another muscle related Module #58 with less enrichment, as well as Module #70 for mitochondrial fatty acid beta-oxidation (OR=25.3, FDR<0.0001). HCM is associated with Modules #85, #92, #39, #70, and another mitochondrial Module #2 (OR=16.5, FDR<0.0001), consistent with that cardiomyopathy could result from mitochondrial dysfunction (37). Notably, contained within Module #85, #92 and #39 are 42 genes with the MP *enlarged heart*, including 28 as novel potential candidate genes for cardiomyopathy (Figure 2C and Figure S5).

Other examples of tissue-specific disease modules include those for ciliary motility disorders (#1 and #67), pulmonary alveolar proteinosis (#154), akinesia (#56), hereditary pancreatitis (#105), congenital hypothyroidism (#51), and night blindness (#114) (Table S4).

### Gene modules associated with hypertension

Also identified were modules for hypertension, a complex and multifactorial disease that affects nearly one billion people worldwide (38). Using curated gene-disease associations from DisGeNET, 13 hypertension modules were identified (Table 2 and Table S4), 7 of which form two large categories. The first category are 2 modules regulating sodium or ion homeostasis. Module #281, expressed specifically in kidneys (Figure S6), is enriched with 10 *ion transmembrane transporter* genes (*P*=1.02E-06). It contains 5 hypertension genes (OR=16.5, FDR=0.0003) (Figure 3A), including *UMOD* that encodes uromodulin, the major secreted protein in normal urine. Noncoding risk variants increase *UMOD* expression, which causes abnormal activation of renal sodium cotransporter SLC12A1 and leads to hypertension (39). Interestingly, *SLC12A1* is also in Module #281, together with *SLC22A2* and *TMEM72*. All three genes were identified as blood pressure loci in a GWAS study based on genetic analysis of over 1 million people (40), but not in DisGeNET. The module contains additional genes with the MP *abnormal kidney physiology*, among which might include novel candidate genes for hypertension, as kidneys are important for blood pressure control (41). Module #305, expressed more broadly than #281 (Figure S6), is enriched with 4 hypertension genes (OR=17.6, FDR=0.0016) like *SCNN1B* and *SCNN1G*, which also modulate renal sodium homeostasis (42).

**Figure 3.**
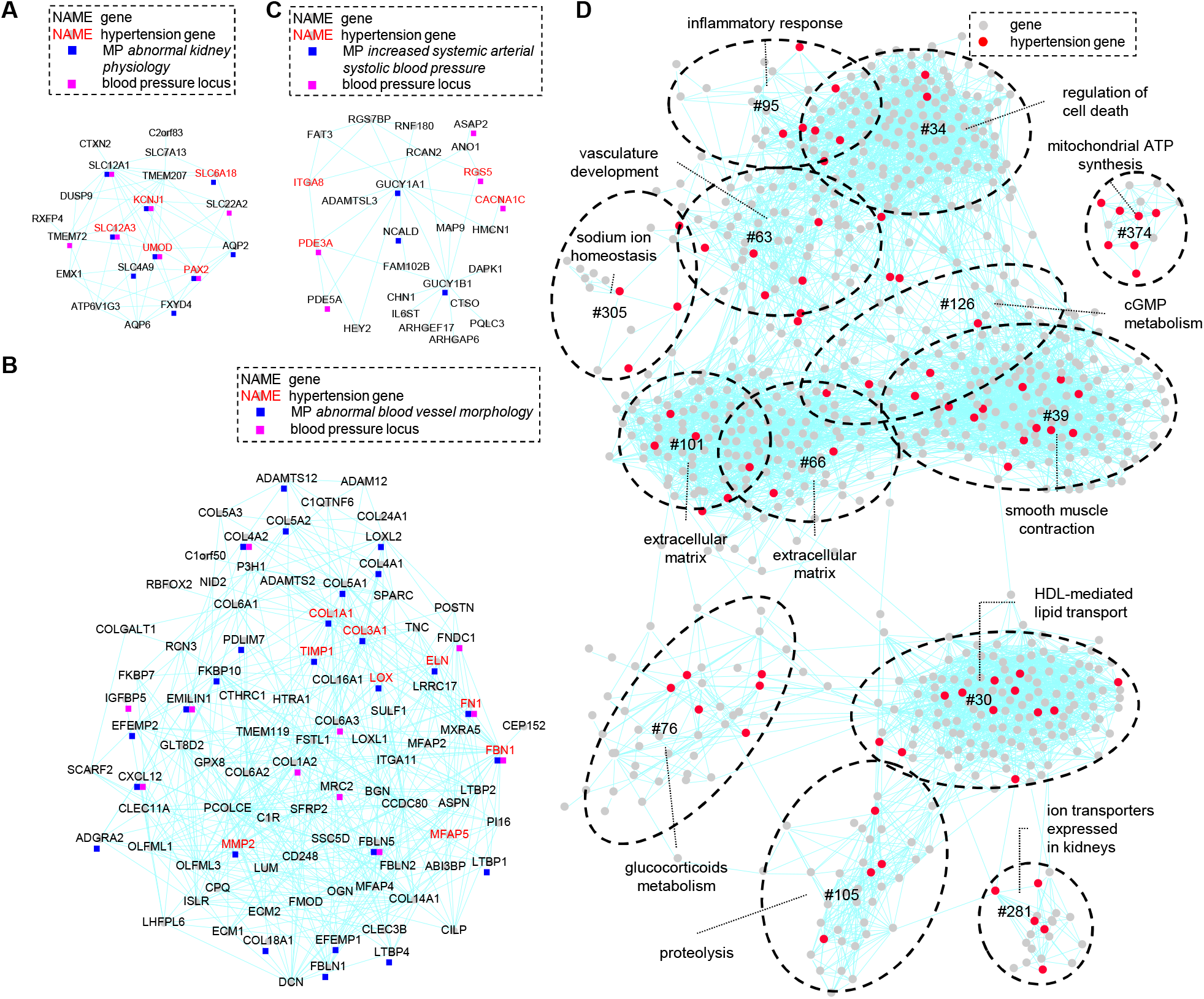
Gene modules associated with hypertension. **(A-C)** Modules #281, #66, and #126 associated with hypertension, respectively. **(D)** A sub-network including all hypertension modules. Gene names are not labeled due to space limitation. Dashed circles outline the approximate positions of the modules.

The second category are 5 modules involved in vascular tone regulation. Module #66, over-represented with *extracellular matrix* genes (*P*=1.59E-75), contains 9 hypertension genes (OR=6.5, FDR=0.0004), such as *ELN* and *FBN1* (Figure 3B). Its enriched MPs include *abnormal blood vessel morphology* and *abnormal blood circulation*, indicating that, similar to *ELN* and *FBN1* (43), genes in the module could determine vascular extracellular matrix composition and regulate arterial compliance. Similarly, Module #101 is also enriched with 40 *extracellular matrix* genes as well as 6 hypertension genes (OR=4.5, FDR=0.0421). Module #39, functioning in smooth muscle contraction, encompasses 13 hypertension genes (OR=5.5, FDR<0.0001), among which are *KCNMB1* and *PRKG1*, encoding key modulators of smooth muscle tone (44,45). Genes within the module, enriched with MPs *abnormal vascular smooth muscle physiology* and *impaired smooth muscle contractility*, could regulate arterial wall elasticity. Module #54, for glucocorticoids metabolism, contains 6 hypertension genes (OR=7.4, FDR=0.0026), and #126 (Figure 3C), participating in cGMP metabolism, includes 4 hypertension genes (OR=10.1, FDR=0.0147). Glucocorticoids and cGMP modulate blood pressure via regulating peripheral vascular resistance and blood vessel relaxation respectively (46,47). The relevance of these 5 modules are further supported by that, besides the hypertension genes in DisGeNET, Module #39, #54, #66, #101, and #126 contain 14, 3, 9, 9, and 2 additional blood pressure loci respectively (Figure 3B, C, and Figure S7), as revealed by the above mentioned GWAS study (40).

The rest 6 hypertension modules function in inflammatory response (#95), HDL-mediated lipid transport (#30), mitochondrial ATP synthesis (#374), regulation of cell death (#34), vasculature development (#63), and proteolysis (#105), respectively. Figure 3D summarizes the modules associated with hypertension, which reveal the pathways underlying the pathogenesis of hypertension.

### Gene modules associated with autism

Our approach also identified modules associated with brain disorders, among which is autism, a development disorder characterized by impaired social interaction and by restricted and repetitive behaviors (48). Using SFARI autism genes (5), 12 autism modules were detected (Table 2 and Table S4), 8 of which form three large categories. The first category is a module involved in *covalent chromatin modification* (#7), which contains 221 genes, including 48 autism genes (OR=5.9, FDR<0.0001) (Figure 4A), such as *ASH1L*, *CHD8*, and *KMT2C* (49). The module has 40 genes with the MP *abnormal brain morphology*, among which 12 are autism genes. It is also over-represented with MPs *abnormal heart morphology*, *abnormal gastrulation*, and *abnormal axial skeleton morphology*. Thus gene mutations in this module might affect multiple developmental processes, including brain development that could lead to autism.

**Figure 4.**
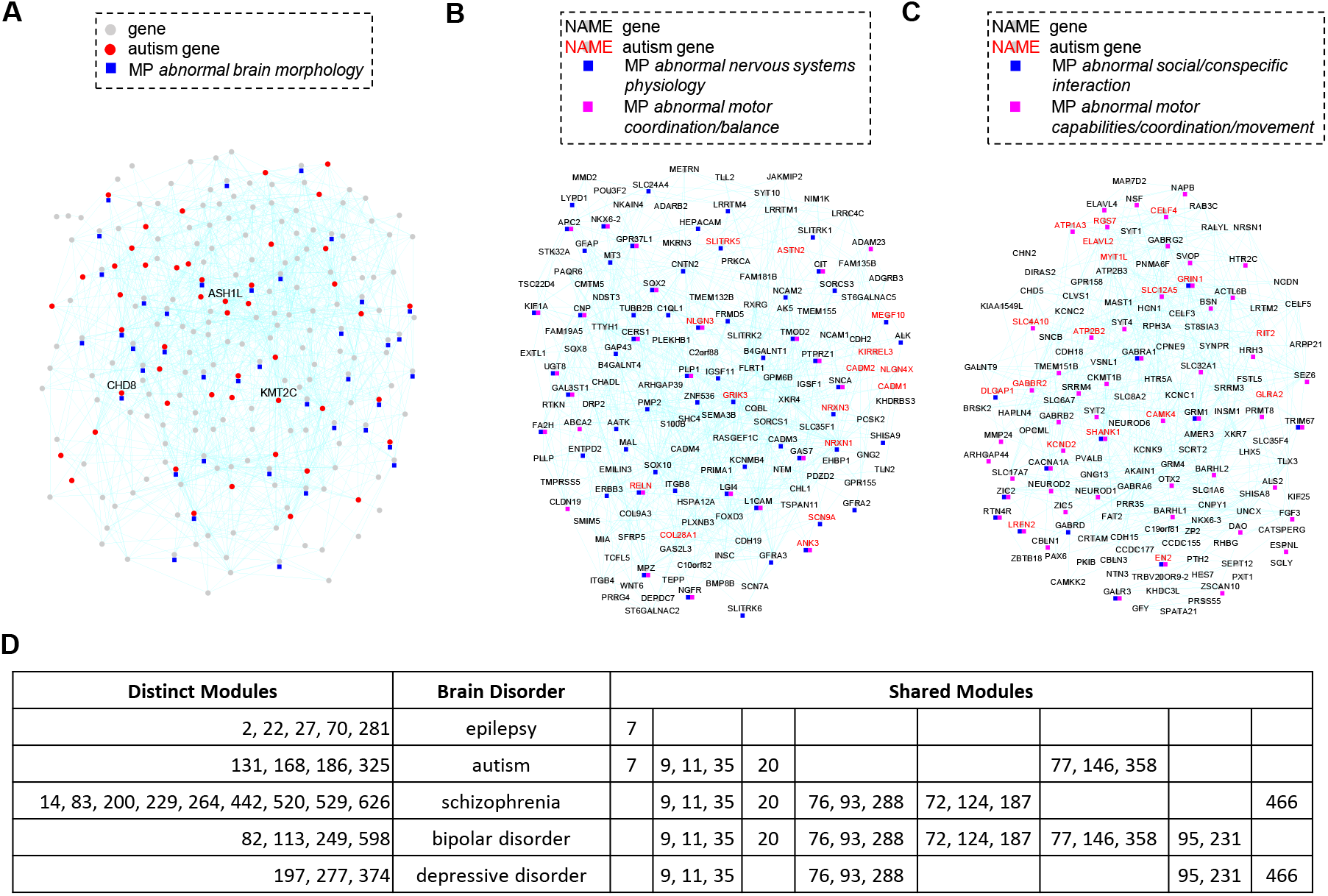
Gene modules associated with autism and other brain disorders. **(A-C)** Modules #7, #9, and #20 associated with autism, respectively. Most gene names are not labeled in (A) due to space limitation. **(D)** Distinct and shared disease modules among epilepsy, autism, schizophrenia, bipolar disorder, and depressive disorder. Numbers indicate the module ids.

The second category are 2 modules involved in *nervous system development* (#77) and *neurogenesis* (#9). Among them, Module #9 contains 15 autism genes (OR=2.5, FDR=0.0356) (Figure 4B), like *NLGN3*, *NLGN4X*, and *NRXN1* (50). The module is enriched with autism-related MPs, such as *abnormal nervous system physiology* and *abnormal motor coordination/balance* (51). Among the 60 genes with the MP *abnormal nervous system physiology*, 9 are autism genes and the rest might contain additional candidate genes.

The third category are 4 modules participating in synaptic signaling. Though all enriched with *synaptic signaling* genes (*P*=3.66E-36 to 7.33E-10), these modules express differently. Module #20 and #35 have biased expression in cerebellum and cortex respectively, while #11 and #146 express broadly in various brain regions (Figure S8). All 4 modules are enriched with autism genes and might contain potential candidate genes. For example, Module #20 has 18 autism genes (OR=3.3, FDR=0.0006) (Figure 4C), like *MYTL1* and *DLGAP1* (16,49). It also contains 55 genes with MPs *abnormal social/conspecific interaction* and/or *abnormal motor capabilities/coordination/movement*, among which are 14 autism genes and the rest merit further investigation.

Among the other autism modules, #186 and #131 function in cell-cell adhesion and extracellular matrix organization respectively, #358 is enriched with *neuron part* genes, while #168 and #325 are without clear biological interpretation presently. Thus, these modules indicate abnormality in chromatin modification, nervous system development, synaptic signaling and other processes could lead to autism.

### Shared and distinct disease gene modules among brain disorders

Modules for epilepsy, schizophrenia, bipolar disorder, and depressive disorder are also identified, revealing shared and distinct disease modules among brain disorders (Figure 4D and Table S4). Module #7 for chromatin modification is associated with autism and epilepsy only, but Modules #11 and #35 for synaptic signaling and #9 for neurogenesis are shared by autism, schizophrenia, bipolar disorder and depressive disorder. Also shared by bipolar disorder and depressive disorder are Module #231 involved in circadian rhythm regulation and Module #95 for inflammatory response, highlighting the roles of circadian rhythm and inflammation in mood disorders (52–54). On the other hand, epilepsy is distinctly associated with modules functioning in mitochondria (#2, #22, #70), lysosomes (#27), and kidneys (#281), indicating its unique etiology (55–57), while schizophrenia is uniquely associated with modules for MHC antigen presentation (#83, #264), myelination (#14), and fatty acid biosynthesis (#520) (58,59). The rest shared or distinct brain disorder modules also function in synaptic signaling (#93, #146, #187, #229), ion transport (#72, #113), neuropeptide signaling (#277), and dopamine metabolism (#288). These modules delineate the pathways associated with brain disorders and should facilitate their etiology studies.

## DISCUSSION

Our GGM co-expression network analysis identified unbiased data-driven gene modules with enriched functions in a variety of pathways and tissues. Besides modules with tissue-specific expression, such as those for cilium (Module #1), myelin sheath (#14), fat (#18), liver (#30), brush border (#50), and thyroid (#51), it also recovered modules co-expressed broadly in multiple tissues, i.e. those for cholesterol biosynthesis (#71), protein folding (#29), and mitochondria or lysosome functions (#2, #22, #27). It distinguished modules with closely related functions, such as the two modules regulating sodium homeostasis with kidney specific expression (#281) and broad expression patterns (#305), the modules for striated muscle (#85 and #92) and smooth muscle (#39) functions, and the modules regulating vascular tone via extracellular matrix composition (#66 and #101) and smooth muscle contraction (#39) respectively. The high resolution power and broad coverage of our GGM network could be attributed to the usage of partial correlation coefficient for network construction, which measured correlation between two genes while controlling for the effect of other genes and thus only direct correlated gene pairs were retained (19,20). It could also be due to that a large amount of GTEx samples from 53 tissue types were used in the analysis, enabling the network to uncover the co-expression modules underlying all these tissues. Additionally, the identified modules were not too large in size, containing between 9 to 354 genes each, which laid a solid foundation for subsequent disease modules identification. Note that the modules could also be used to study normal physiological pathways under healthy conditions.

Although the GGM network was derived from non-diseased samples, disease genes converge on the identified gene modules. Such convergence was expected for diseases affecting specific tissues or organs. For example, many cardiomyopathy disease genes are enriched in the modules for both started and smooth muscle functions (#85, #92, #39), hereditary pancreatitis genes are preferably mapped to a pancreas specific module (#105), while congenital hypothyroidism are concentrated in a thyroid specific module (#51). Convergence was also observed for complex and multifactorial diseases like obesity, hypertension, and autism. Multiple diseases gene modules were identified for these diseases, reflecting the heterogeneous nature of their pathogenesis. The disease gene modules could represent the molecular pathways underlying different branches of diseases’ etiologies. Our analysis provided a general summary of the modules for, i.e., obesity (Figure 2B), hypertension (Figure 3D), autism, and other brain disorders (Table S4). Comparison between diseases began to reveal shared and distinct gene modules, which was not limited to brain disorders but also expanded to others. For example, Module #39 for smooth muscle contraction is associated with both cardiomyopathy and hypertension, while Module #277 for neuropeptide pathway is involved in both obesity and depressive disorder. Such comparison will further enhance our understanding of disease mechanisms. Additionally, the identified disease gene modules, mostly with clear biological interpretation, integrate well with previous disease knowledge. We believe such modules will facilitate disease etiology studies in the future.

Besides GO and disease genes enrichment, the identified disease gene modules were further supported by large-scaled mouse model phenotypes recoded as MP assignments. Different from GO assignments that may be inferred from electronic annotation, MP assignments were based on the phenotype examination of actual null or transgenic mouse gene models (30). We passed mouse genes’ MP assignments from MGI onto their human orthologues, and found many related MPs were enriched within the identified diseases gene modules, as discussed above. Such MP enrichment validated our diseases gene modules and pinpointed potential novel disease candidate genes. Thus, our network and modules also provide a general framework to analyze large-scale mouse models’ phenotypes. Such analysis could also be extended to mouse gene network studies.

In concussion, our GGM gene network analysis identified gene modules associated with multiple diseases as discussed above. It also works for many other human diseases with genetic basis. The identified module can be used to pinpoint the pathways underlying disease pathogenesis and identify potential novel disease genes. It merits investigating whether it can also be used to categorize disease subtypes to guide treatment development accordingly. Future analysis can also include non-coding genes, such as long non-coding RNAs, to study their roles on diseases development.

## MATERIALS AND METHODS

### Co-expression network construction

Open-access and de-identified GTEx V7 transcriptome data were downloaded from the GTEx Portal (http://www.gtexportal.org). The data were fully processed and available as a matrix that contains TPM (Transcripts Per Million) gene expression values for 53035 genes in 11688 samples spanning 53 tissues from 714 postmortem donors. After filtering out low expressed genes that have TPM values >=1 in less than 10 samples, the expression data were normalized in a tissue-aware manner via qsmooth with default parameters (60). A sub-matrix consisting of 18626 protein-coding genes was extracted from the normalized expression matrix and used for GGM network construction, following a procedure described previously (23,24). Briefly, the procedure consisted of 6000 iterations. In each iteration, 2000 genes were randomly selected and used for partial correlation coefficient (pcor) calculation via the GeneNet package (v 1.2.13) in R (19). After 6000 iterations, every gene pair was sampled in average 69 times with 69 pcors calculated, and the pcor with lowest absolute value was selected as its final pcor. Also calculated were Pearson’s correlation coefficient (r) between gene pairs. Finally, gene pairs with pcor>=0.035 and r>=0.35 were chosen for network construction.

### Network clustering and module analysis

The network was clustered via the MCL clustering algorithm with parameters “-I 1.55–scheme 7” (25). The identified modules with >=9 genes were kept, and they were further expanded by including outside genes that connect with >= 4 genes within the original modules. The sub-networks for the modules were visualized in Cytoscape v 3.40 (61). GO enrichment analysis were performed via hypergeometric test, with GO annotations retrieved from Ensembl BioMart (https://www.ensembl.org/biomart) on 03/06/2019. Mouse Mammalian Phenotype (MP) term assignments were obtained from the MGI database (http://www.informatics.jax.org/downloads/reports/MGI_GenePheno.rpt) on 03/07/2019. MP term assignments derived from mouse models involving 2 or more genes were excluded. Mouse genes’ MP assignments were passed on to their human orthologues and used for MP enrichment analysis via hypergeometric test.

### Identification of modules associated with diseases

Gene-disease associations for obesity were obtained from the human obesity gene map (4). Autism genes were obtained from the SFARI database on 01/15/2019, and only the autism genes scoring as category 1 to category 4 were used in the analysis (5). For other diseases, the curated gene-disease associations registered in DisGeNET v5.0 were used (3). For hypertension, besides the disease genes from DisGeNET, blood pressure loci were also obtained from a recent GWAS study, by combining the previous reported blood pressure loci and the newly confirmed variants, as listed in Supplementary Table 4 and 5 in the article by Evangelou *et al.* (40).

For every disease, its disease gene list were queried against every gene module to calculate a pValue for disease gene enrichment using hypergeometric test. Suppose a disease has *m* disease genes within a gene module with the size of *k*, and *M* disease genes among all *K* genes in the whole network. A pValue for that disease and module combination was calculated as:

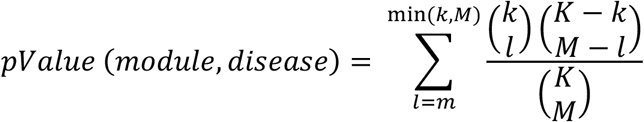

For every disease, the pValues for all modules were adjusted for multiple testing via the Benjamini-Hochberg procedure (62).

A permutation based procedure was also used to estimate false discovery rate (FDR). In each permutation, every disease’s disease gene list were replaced by the same number of genes randomly selected from the whole network and used for enrichment calculation. After conducting 10000 permutations, the results were tallied and used to calculate the FDRs corresponding to original pValues. Modules with FDR less than or equal to 0.05 and with 3 or more disease genes were kept for further analysis.

## SUPPLEMENTARY MATERIALS

**Table S1:** The 166980 co-expressed gene pairs used for network construction

**Table S2:** The 652 gene modules identified from the network

**Table S3:** GO enrichment analysis results

**Table S4:** Disease genes enrichment analysis results

**Table S5:** MP enrichment analysis results

**Figure S1:** Gene expression heatmap for Modules #18, #30, and #71

**Figure S2:** The pathway for synthesizing cholesterol from acetyl-CoA

**Figure S3:** Gene expression heatmap for Modules #50 and #71

**Figure S4:** Gene expression heatmap for Modules #39, #85, and #92

**Figure S5:** Modules #39 and #92 associated with cardiomyopathy

**Figure S6:** Gene expression heatmap for Modules #281 and #305

**Figure S7:** Modules #39, #54, and #101 associated with hypertension

**Figure S8:** Gene expression heatmap for Modules #11, #20, #35, #76, and #124

## ACKNOWLEDGEMENT

We thank the GTEx donors and their families for their contribution to science and the GTEx consortium for generating the resource. The Genotype-Tissue Expression (GTEx) Project was supported by the Common Fund of the Office of the Director of the National Institutes of Health and by NCI, NHGRI, NHLBI, NIDA, NIMH, and NINDS. The GTEx transcriptome data used for the analyses described in this manuscript were obtained from the GTEx Portal (http://www.gtexportal.org) on 12/05/2017. The work in Ma Lab is supported in part by grants from National Natural Science Foundation of China (31770268), Fundamental Research Funds for the Central Universities (WK2070000091), and University of Science and Technology of China (Start-up fund to S.M.). The numerical calculations in this manuscript were conducted on the supercomputing systems in USTC Supercomputing Center and in USTC School of Life Sciences Bioinformatics Center.

