## Supplemental Figures for "Gene modules associated with human diseases revealed by network analysis"

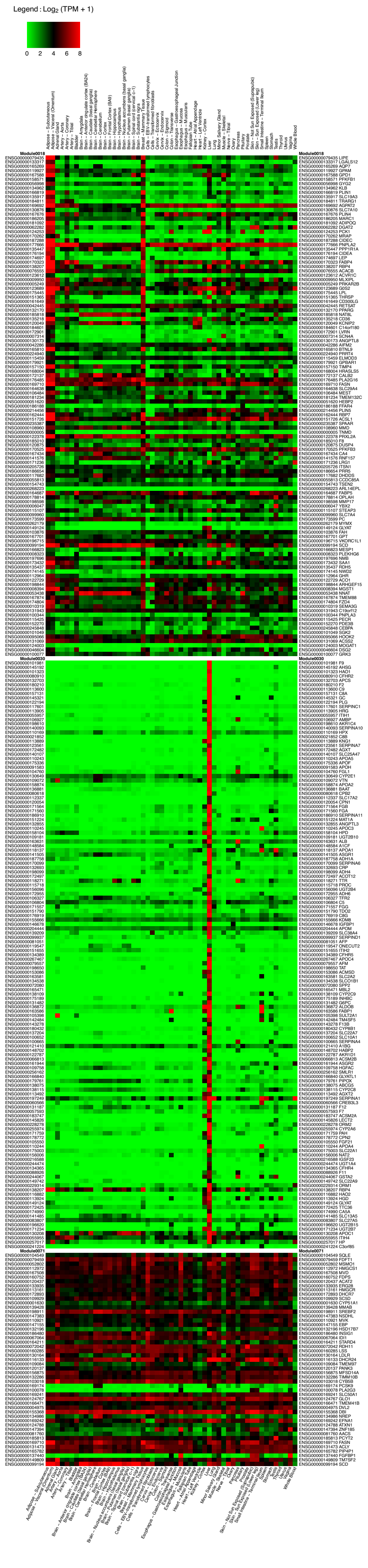


**Figure S1**

**Gene expression heatmap for Modules #18, #30, and #71.** Shown are the median gene expression values by tissues according to GTEx.

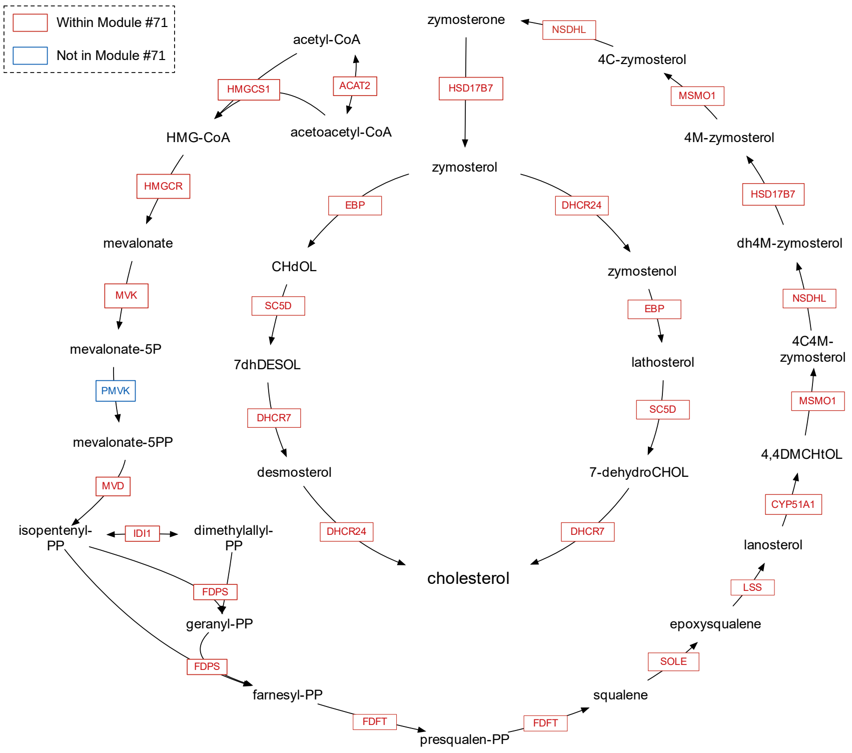


**Figure S2**

**The pathway for synthesizing cholesterol from acetyl-CoA.** Module #71 contains genes encoding enzymes for 23 of all 24 reaction steps of the pathway. Red rectangles indicate the corresponding genes are within Module #71. Blue rectangle indicates the corresponding gene is not in the module. The pathway was obtained from Reactome (https://reactome.org/).

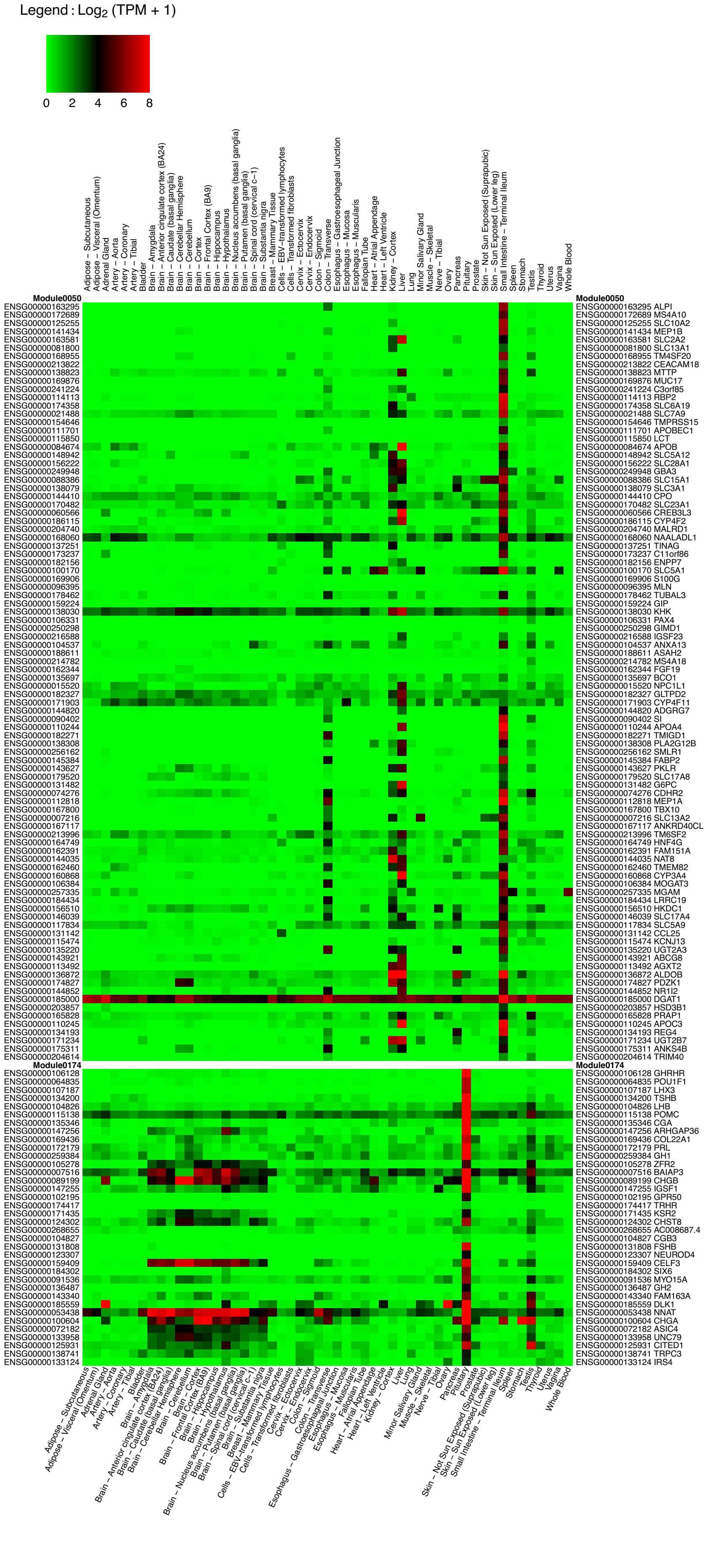

**Figure S3**

**Gene expression heatmap for Modules #50 and #71.** Shown are the median gene expression values by tissues according to GTEx.

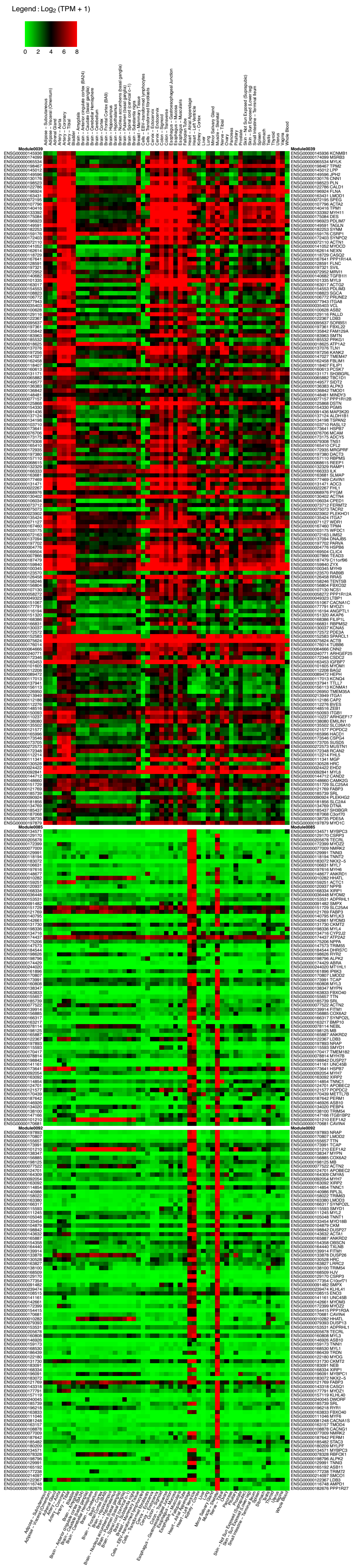

**Figure S4**
**Gene expression heatmap for Modules #39, #85, and #92.** Shown are the median gene expression values by tissues according to GTEx.

A

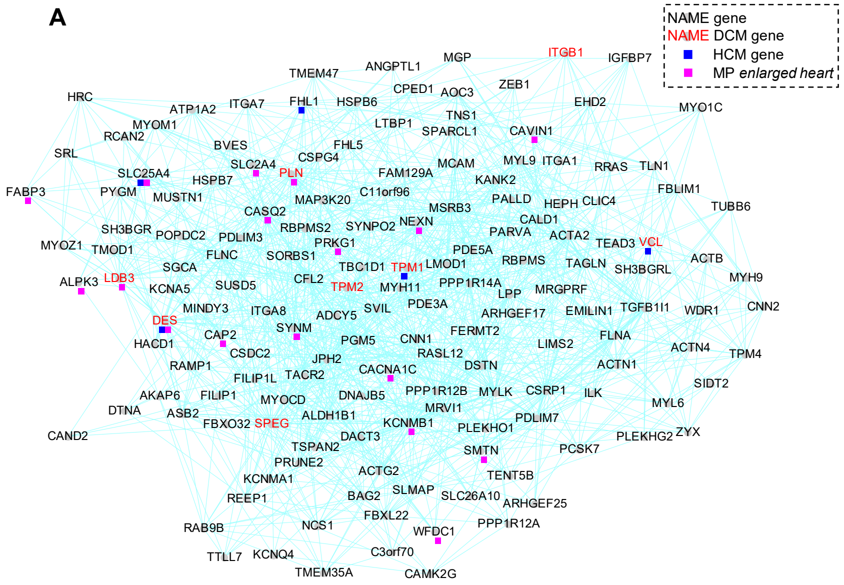


B

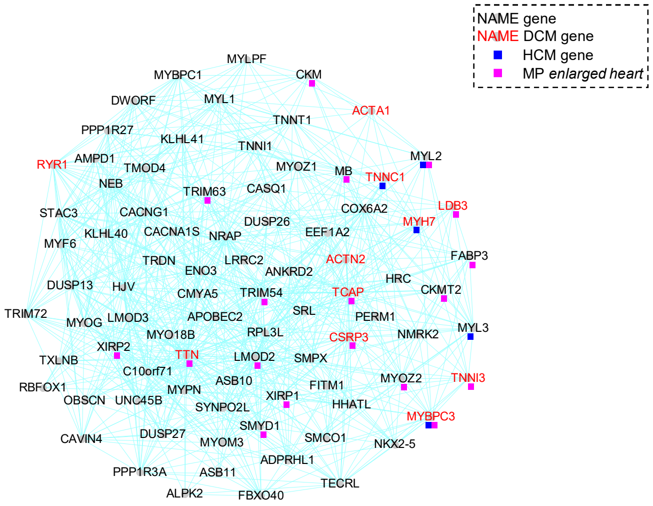


### Figure S5
### Modules #39 (A) and #92 (B) associated with cardiomyopathy.

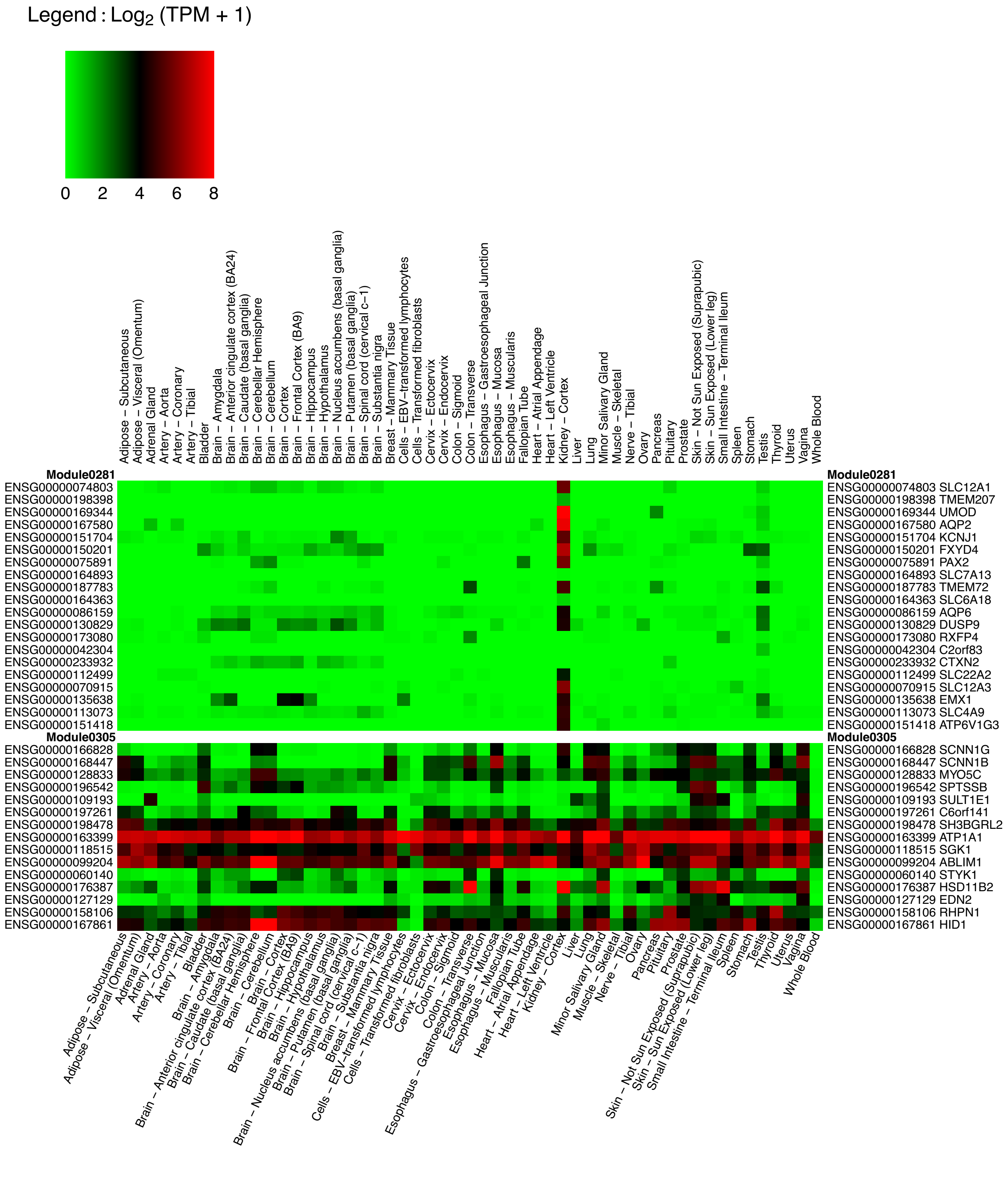


**Figure S6**

**Gene expression heatmap for Modules #281 and #305.** Shown are the median gene expression values by tissues according to GTEx.

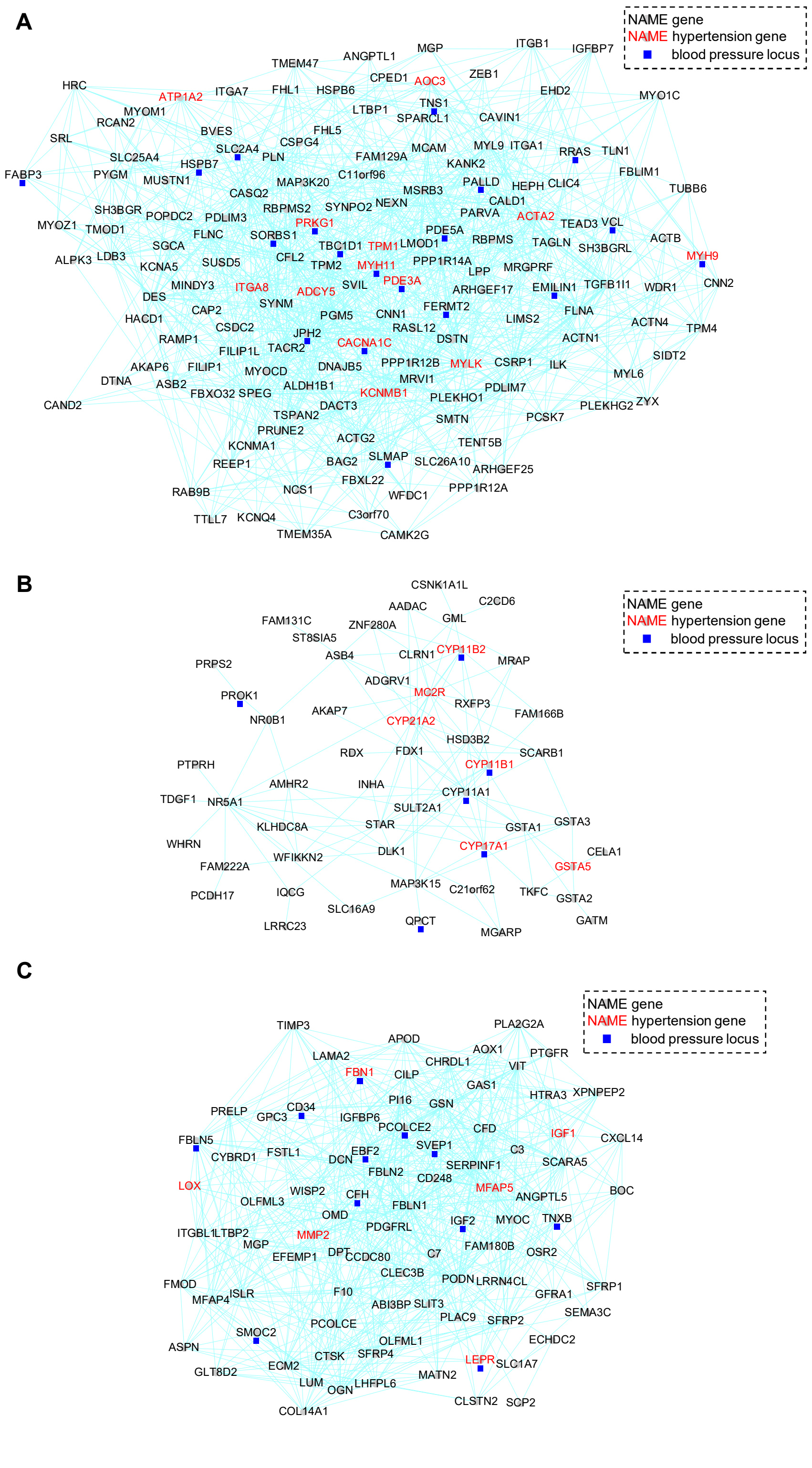


**Figure S7**

**Modules #39 (A), #54 (B), and #101 (C) associated with hypertension.**

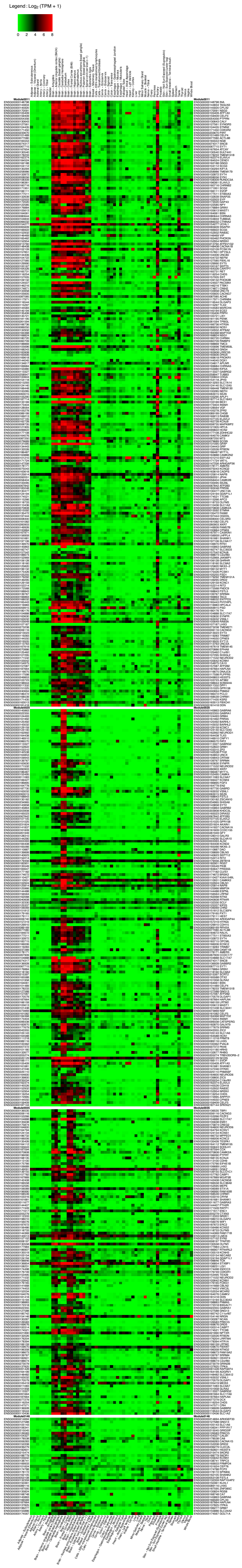

**Figure S8**

**Gene expression heatmap for Modules #11, #20, #35, and #146.** Shown are the median gene expression values by tissues according to GTEx.
